# From Code to Cure: Computationally Designed BMP-2 Binders Using AI-Integrated Pipelines for Controlled Bone Regeneration

**DOI:** 10.64898/2026.08.20.745793

**Authors:** Ben Burress, Arash Asgari, Jonathan Dorogin, Karly Fear, Cassandra Gonzalez, Justin Svendsen, Dylan Merrill, Marian H. Hettiaratchi, Parisa Hosseinzadeh

## Abstract

Nonunion fractures remain a costly and persistent challenge in regenerative medicine, with current treatments limited by donor-site morbidity, restricted graft availability, and severe adverse effects associated with supraphysiological bone morphogenetic protein 2 (BMP-2) delivery, including ectopic ossification and inflammation. Endogenous BMP-2 signaling is tightly regulated in adult tissues, constraining the precision and scalability of approaches based on transcriptional upregulation or bolus growth factor administration. To address these limitations, we developed a two-phase integrated computational–experimental pipeline for the de novo design of protein binders targeting the BMP-2 knuckle epitope, a receptor-binding surface corresponding to BMPR-II engagement, enabling affinity-tuned modulation of BMP-2 activity rather than uncontrolled pathway activation. Phase I employed PyRosetta-based β-strand motif grafting and physics-based docking protocols to generate 264 candidate binders, followed by deep-learning-driven refinement in Phase II using partial RFDiffusion and ProteinMPNN with AlphaFold2 validation, yielding 22 candidates with stable β-sheet architectures consistent with knuckle-epitope targeting. Experimental validation demonstrated dose-dependent BMP-2 binding, with the lead construct exhibiting an apparent equilibrium dissociation constant of 2.07 nM toward BMP-2. Targeted alanine substitutions revealed differential residue contributions, with mutation of T42 significantly disrupting binding, while other substitutions had more modest effects, indicating a partially hotspot-driven interface supported by other interactions.

## Introduction

Bone fractures are among the most common traumatic injuries worldwide, and approximately 2% progress to nonunion, a condition characterized by stalled healing and failure to restore structural integrity^1,2^. Nonunion fractures impose substantial socioeconomic and healthcare burdens, often resulting in chronic pain, loss of function, and diminished quality of life^3^ Autologous bone grafting remains the clinical gold standard for treatment^4^; however, its widespread use is constrained by donor-site morbidity, finite tissue availability, and variable clinical outcomes, underscoring the need for more precise and effective therapeutic strategies.

Bone morphogenetic protein-2 (BMP-2) is a growth factor that induces osteogenic cell differentiation and promotes bone formation ^5,6^. Its clinical introduction marked a major advance in regenerative medicine, enabling bone formation in otherwise recalcitrant settings. Despite this promise, therapeutic application of BMP-2 is limited by the need for supraphysiological dosing to achieve effective bone formation. High-dose BMP-2 administration has been associated with serious adverse effects, including ectopic ossification, inflammatory responses, and uncontrolled bone growth, which have restricted broader clinical adoption^7^. These challenges highlight the need for strategies that enable localized, sustained, and dose-controlled modulation of BMP-2 activity, within a narrow therapeutic window^8^

At the molecular level, BMP-2 signaling is tightly regulated through interactions with cell-surface receptors and extracellular antagonists. BMP-2 engages two classes of receptors, BMPR-I and BMPR-II, via distinct epitopes known as the wrist and knuckle regions, respectively (Fig. 1).^9^ Receptor assembly triggers activation of the canonical SMAD signaling cascade, ultimately leading to transcription of osteogenic genes^10^. Extracellular inhibitors such as Noggin further modulate BMP-2 availability by sequestering the ligand and preventing receptor engagement. Together, these mechanisms tightly constrain endogenous BMP-2 activity, particularly in adult tissues, limiting the effectiveness of strategies based on transcriptional upregulation or bulk protein delivery^9^

**Figure 1.**
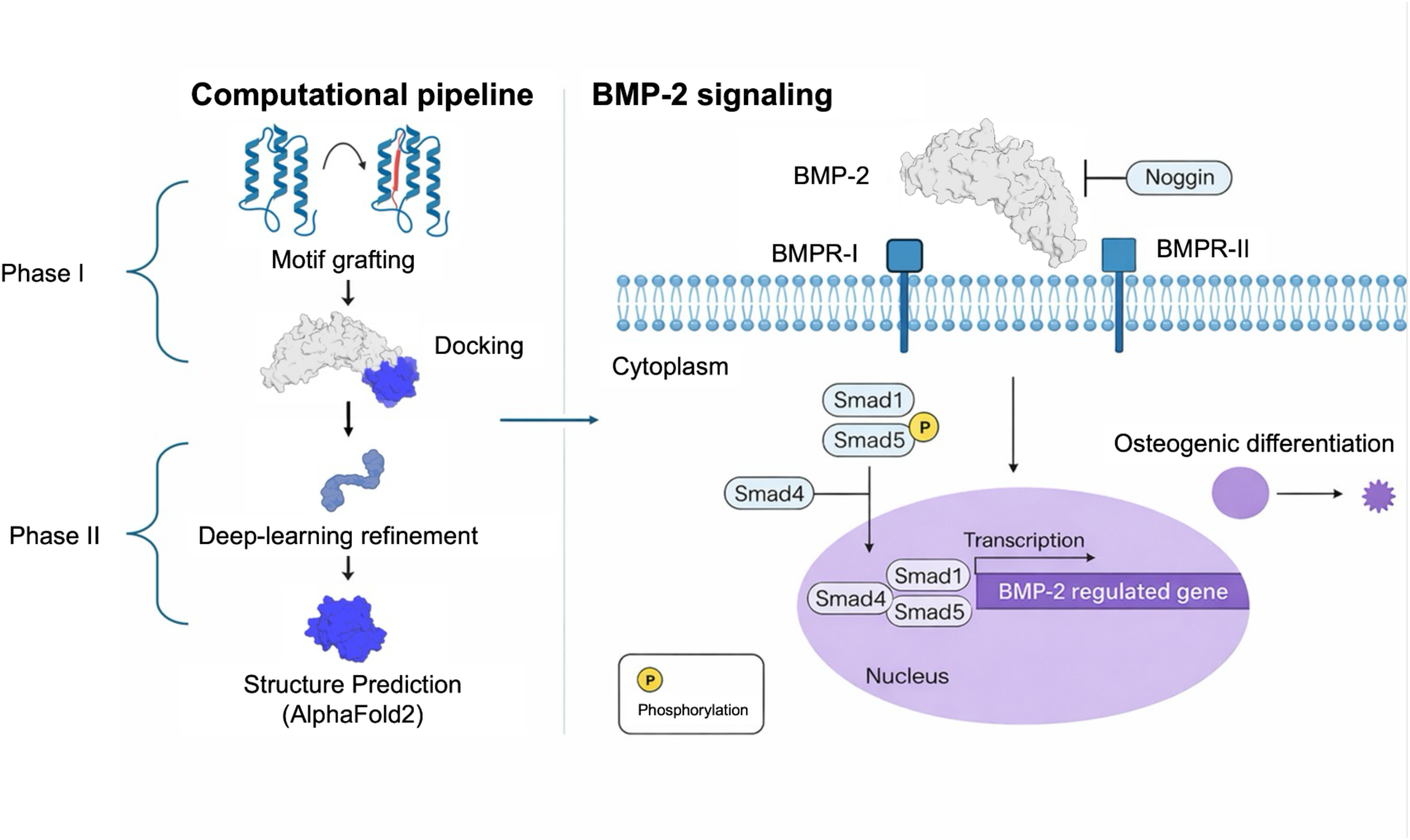
Two-phase computational design workflow and BMP-2–mediated canonical SMAD signaling. Two-phase computational workflow for BMP-2 binder design. Phase I implements Rosetta-based motif grafting and docking to generate initial binder candidates. Phase II applies deep-learning-driven refinement using RFDiffusion and ProteinMPNN, followed by structural validation with AlphaFold2. Selected designs were carried forward for downstream experimental characterization.

An emerging alternative strategy is the use of engineered protein binders to modulate growth factor availability at the protein level. By selectively targeting epitopes critical for receptor engagement, such binders can tune ligand presentation, release kinetics, and signaling output without requiring supraphysiological dosing^11,12^ In this context, the BMP-2 knuckle epitope represents an attractive yet challenging target. This region is structurally complex, β-sheet–rich, and directly involved in receptor binding, making it well suited for β-strand–mediated molecular recognition, yet difficult to target using conventional design approaches that bias toward α-helical bundles^13,14^

Here, we present an integrated computational–experimental platform for the de novo design of protein binders targeting the BMP-2 knuckle epitope. The design strategy was implemented in two distinct phases. Phase I employed PyRosetta-based β-strand motif grafting and Rosetta-based docking protocols to generate an initial library of candidate binders^15–23^. Although representative Phase I designs exhibited favorable Rosetta total-energy scores^24^ and docking metrics, they failed to demonstrate measurable binding to BMP-2 experimentally (SI Fig. 1), highlighting the limitations of physics-based modeling alone for this epitope. These findings motivated Phase II, which leveraged deep-learning–driven backbone refinement and sequence optimization to improve structural accuracy, interface complementarity, and binding fidelity. Phase II also incorporated modern structure-prediction approaches for evaluating protein-protein complexes^25^

Through iterative computational refinement and experimental validation, this two-phase strategy yielded high-affinity BMP-2–specific binders with tunable binding properties (Fig. 1). By modulating growth factor availability at the protein level, this approach offers a potential alternative to bolus BMP-2 delivery and provides a generalizable framework for designing precision binders targeting complex protein–protein interfaces. More broadly, this work illustrates the power of combining physics-based design, deep learning, and empirical feedback to address challenges in protein therapeutic development for regenerative medicine applications.

## Results

### Computational Design, Screening, and Phase I Evaluation

Phase I design focused on generating β-sheet–based binder scaffolds tailored to the BMP-2 knuckle epitope^14,23,26^. Using PyRosetta-based motif grafting^16^, we generated an initial library of 264 β-strand–grafted candidate binders (Fig. 2A). These designs were engineered to present complementary β-strand geometries capable of engaging the β-sheet–rich knuckle region critical for BMP-2 interaction with BMPR-II.

**Figure 2.**
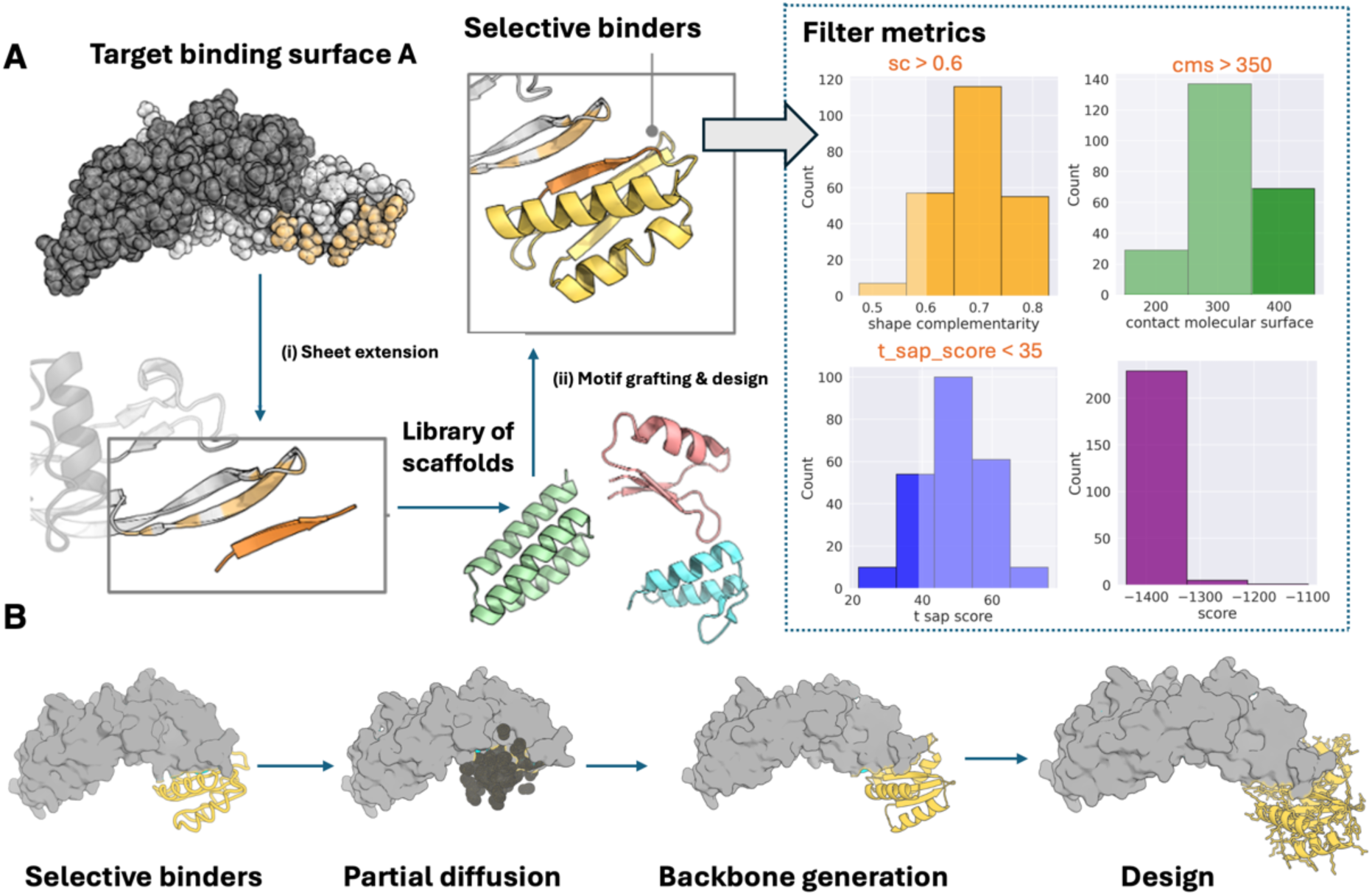
Two-phase computational workflow for generating BMP-2 binder candidates. (A) <u>Phase I:</u> Rosetta-based binder generation. Target binding surface A (the BMP-2 knuckle epitope) was selected as the design site for this campaign. The Phase I workflow consisted of (i) de novo β-sheet extension to generate interface motifs complementary to the knuckle epitope and (ii) grafting of these motifs onto a library of stable scaffold backbones, followed by Rosetta-based design and docking to generate an initial library of candidate binders. (B) <u>Phase I</u> Deep-learning–driven refinement and sequence optimization. Phase I designs were used as inputs to a partial RFDiffusion refinement pipeline. Specifically, (i) Rosetta-designed binders were provided as structural inputs to the deep-learning model, (ii) partial denoising was applied to enable localized backbone reorganization while preserving the overall fold, (iii) refined backbone conformations were generated, and (iv) optimal amino acid sequences were assigned using ProteinMPNN followed by Rosetta FastRelax to improve packing, interface complementarity, and overall structural stability.

Initial prescreening eliminated nonviable constructs based on energetic stability, steric compatibility, and backbone geometry. Designs exhibiting unfavorable Rosetta total energy scores, steric clashes, or disrupted β-strand integrity were excluded. Sequences containing cysteine residues were also removed to minimize the risk of unintended disulfide bond formation and aggregation. This filtering reduced the design pool to eight candidates that met all energetic and steric thresholds.

The remaining candidates were subjected to global docking simulations with BMP-2 using Rosetta^15^. Five designs exhibited convergent, funnel-shaped binding energy landscapes, characterized by improved interface energy scores with decreasing RMSD toward the designed pose (Fig. 3A,B). Such funnels indicate repeated sampling of similar low-energy docking poses rather than isolated low-energy outliers. Additional Phase I docking landscapes are provided in SI Fig. 2. AlphaFold2 predicted backbone RMSDs below the 3 Å filtering threshold relative to the corresponding design models and predicted Local Distance Difference Test (pLDDT) scores exceeding 80 across the structured core, indicating strong agreement between the predicted and designed structures (Fig. 4A,B).

**Figure 3.**
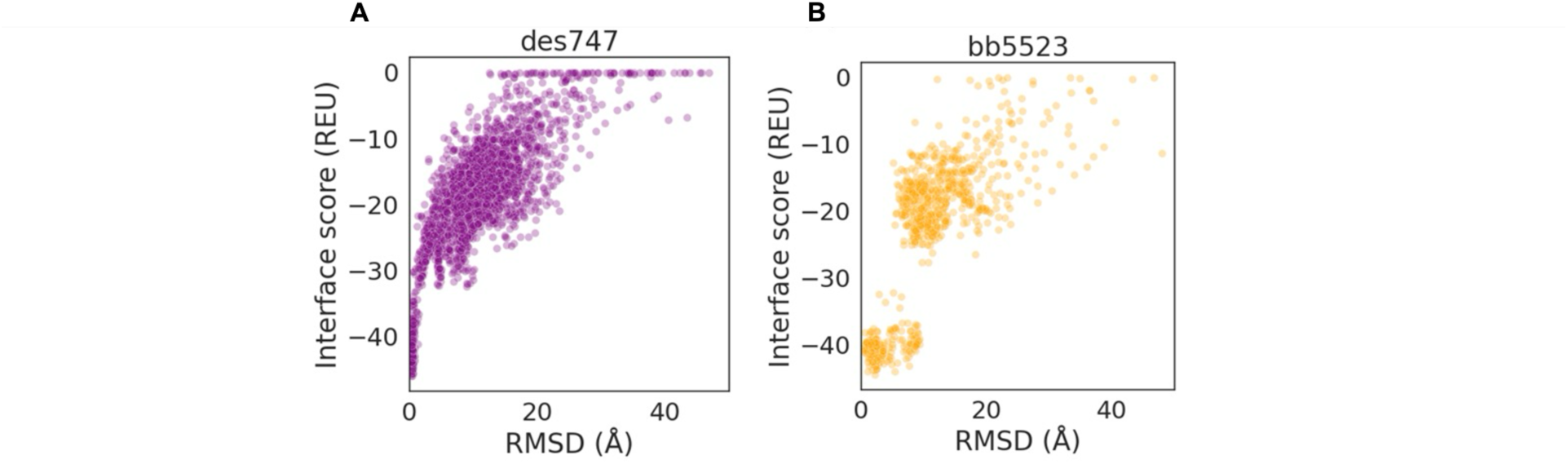
Docking simulations reveal funnel-shaped binding energy landscapes for BMP-2 binding. Each point represents a docked decoy complex generated by RosettaDock, plotted as interface energy score (I_sc) versus RMSD relative to the designed binding pose. Phase I Rosetta-designed binders (des624, des015, des004, and des747) docked against BMP-2 exhibit funnel-shaped binding landscapes with varying degrees of convergence, ranging from diffuse and weakly populated funnels to more pronounced, smoothly convergent energy minima. Among these, des747 exhibited the most densely populated low-RMSD energy minimum toward the designed pose. In contrast, the deep-learning–refined Phase II BB5523 (orange) does not exhibit a well-defined low-RMSD energy funnel, but it was retained due to stronger alternative metrics from AlphaFold2, suggesting overall structural reliability despite lacking a well-populated low-RMSD docking funnel.

**Figure 4.**
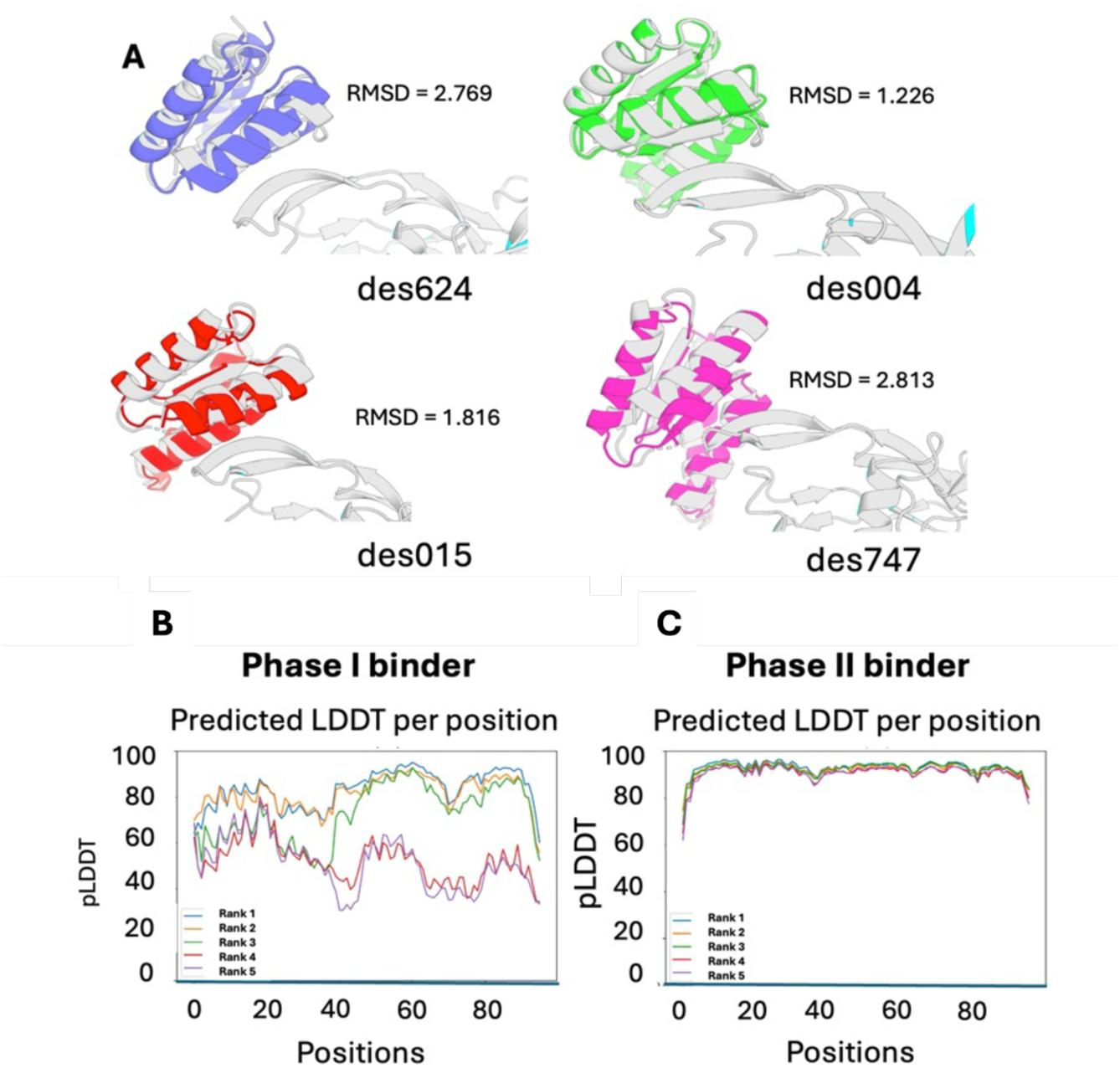
Deep-learning refinement improves structural confidence and model agreement of BMP-2 binders. Structural overlays of AlphaFold2-predicted binder models superimposed on the corresponding Rosetta design models demonstrate improved agreement following Phase II refinement, as reflected by reduced backbone RMSD values (Å) relative to Phase I designs. (A) Structural overlays of AlphaFold2-predicted models and corresponding Rosetta design models (blue = des624, green = des 004, red = des015, pink = des747, gray = BMP-2). for representative Phase I and Phase II binders. (B) Per-residue AlphaFold2 pLDDT scores for the top-ranked Phase I motif grafted binder show variable confidence across the sequence, particularly in flexible and non-core regions. (C) Following Phase II deep-learning–driven refinement, per-residue pLDDT scores are uniformly elevated (>80) across the binder, indicating improved local structural confidence and fold stability. Together, these results demonstrate that RFDiffusion- and ProteinMPNN-based refinement increased pLDDT scores and reduced backbone RMSD between design models and AlphaFold2 predictions.

Of the five computationally prioritized candidates, four were successfully advanced to experimental testing. Despite favorable computational docking behavior, representative Phase I binders expressed experimentally did not exhibit measurable binding to BMP-2 above baseline or negative-control responses (SI Fig. 3). Collectively, these results indicate that favorable docking funnels alone are insufficient predictors of experimental binding for β-sheet–mediated interfaces and motivated the development of a second design phase incorporating deep-learning–driven refinement.

### Phase II: Deep-Learning Refinement, Confidence Filtering, and Docking Validation

To address the limitations observed in Phase I, we initiated Phase II refinement using the highest-rated Phase I scaffold as a structural seed. Using the highest-ranked Phase I scaffold as a starting point, a total of 1,000 partial-diffusion backbone designs were generated and sequence-optimized using ProteinMPNN^27,28^. These designs were subsequently filtered based on Rosetta energetic stability, predicted interface quality and AlphaFold2 confidence metrics, yielding 22 candidates selected for experimental characterization (Fig. 2B)^27^. Collectively, these refinements improved pLDDT scores, reduced interface predicted Aligned Error (pAE) and increased alignment between design and AlphaFold2-predicted structures^29^

After refinement, candidates were filtered using AlphaFold2 confidence metrics. Structural confidence was evaluated using pLDDT (Fig. 4C), a per-residue estimate of interface prediction reliability. Interface quality was assessed using interface predicted aligned error (pAE), which measures confidence in the relative positioning of residues across a predicted interface, to prioritize models with pLDDT > 80 and interface pAE < 10 Å ^29^. Designs that passed confidence thresholds were then evaluated by global docking against BMP-2 to assess preservation of the intended binding mode. AlphaFold-Multimer predictions were additionally used to evaluate binder-target complex geometry and agreement with the designed interface^25^

Although conventional docking metrics did not show corresponding improvement -- and in some cases appeared less favorable -- we advanced refined designs based on improved structural confidence, interface organization, and agreement with AlphaFold2 predictions. This decision reflects the recognition that docking scores alone do not capture the full complexity of protein–protein interactions or reliably predict experimental binding outcomes, particularly for β-sheet–mediated interfaces^30^. Similar observations have been reported in recent de novo binder-design studies in which deep-learning-derived confidence metrics improved experimental success rates relative to energy-based design metrics alone^31^

Following combined filtering and docking evaluation, 22 Phase II candidates were selected for experimental testing, yielding a single BMP-2 binder (BB5523, 1/22 candidates tested) that demonstrated reproducible binding in BLI assays. Notably, the single experimentally validated binder emerged from the subset of designs exhibiting high pLDDT scores, low interface pAE values and agreement between AlphaFold2 predictions and the intended design model, whereas designs selected primarily on docking behavior during Phase I failed to show kinetic binding signal. These findings suggest that modern structure-prediction confidence metrics may provide a more reliable indicator of binder quality than docking scores alone for β-sheet mediated interfaces. Targeted single-point variants of this lead design were subsequently generated to systematically probe the contribution of predicted interface residues, with mutations expected to reduce binding affinity and thereby validate the designed surface.

### Experimental Validation Confirms Functional BMP-2 Binding

To probe the predicted binding interface, targeted mutations were generated using the structural model of the BB5523:BMP-2 complex. The designed interface is mediated by an antiparallel β-sheet interaction between the BB5523 strand FTWIELD and a complementary strand in the BMP-2 knuckle epitope (approximately KVVLKN). Structural analysis suggested that T42, I44 and L46 contribute predicted hydrogen-bonding and van der Waals interactions to the binding interface, whereas F41, W43 and E45 primarily participate in backbone-mediated hydrogen bonding required for β-sheet pairing. Therefore, T42, I44 and L46 were selected for alanine-scanning mutagenesis to assess their energetic contributions to binding. T42 was also mutated to asparagine to distinguish the effects of side-chain truncation from retention of a polar functional group.

The lead Phase II binder (BB5523) and its designed variants (T42A, I44A, L46A, as well as T42N for comparison and T42A/I44A/L46A) were successfully expressed and purified, with all constructs exhibiting molecular masses consistent with their designed sequences, confirming proper expression and molecular integrity (SI Fig. 3).

### Structural Characterization

Circular dichroism (CD) spectroscopy was used to assess secondary structure and folding integrity (SI Fig. 4). Far-UV CD measurements (190-250 nm) exhibited characteristic minima near 208 and 222 nm, consistent with a well-folded protein containing substantial α-helical content. Normalized overlays of six independently measured spectra revealed highly conserved spectral features across all constructs. Secondary-structure deconvolution using BeStSel further supported this observation^32,33^, with all variants exhibiting related secondary-structure compositions and excellent agreement between experimental and fitted spectra (NRMSD < 0.03; Table SI.2). In particular, the parent construct, T42N, I44A and the triple mutant displayed highly similar distributions of α-helical, β-sheet, turn and irregular structural elements. Collectively, these data indicate that BB5523 and its variants remain folded and that the observed changes in binding affinity are unlikely to arise from global protein unfolding.

### Binding Affinity and Specificity

Binding interactions between BMP-2 and the designed binders were evaluated using biolayer interferometry (BLI) across a concentration series (Fig. 5A–C). The parent construct (BB5523) exhibited strong, concentration-dependent binding consistent with an apparent binding affinity of 2.07 nM. (Fig. 5.).

**Figure 5.**
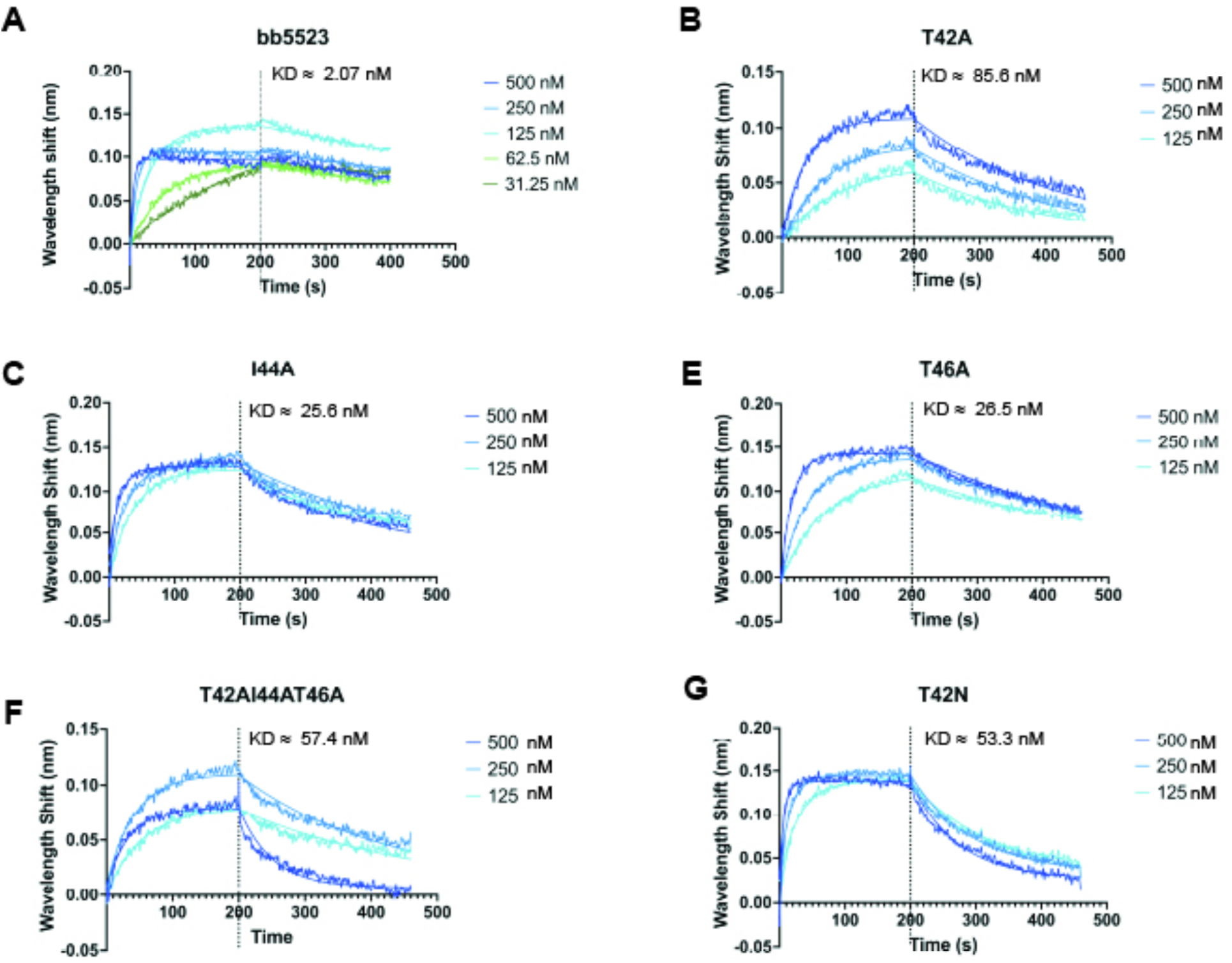
Biolayer interferometry reveals high-affinity and tunable binding of designed BMP-2 binders. (A–C) BLI sensorgrams of the parent construct (BB5523) and mutational variants. Global 1:1 Langmuir kinetic fits across concentration series were statistically rejected (p < 0.0001) due to non-ideal binding behavior. (D) Apparent KD values derived from independent 1:1 kinetic fits localized to the 250 nM concentration curves to enable consistent comparison across constructs.

Statistical comparison of kinetic fits across concentrations (P < 0.0001) indicated that a single global binding model did not adequately describe the data. Consequently, kinetic parameters are interpreted as apparent values useful for comparing variants rather than absolute thermodynamic constants. Possible contributors include heterogeneous binding behavior, surface immobilization effects, or multivalent engagement of the BMP-2 dimer.

The 250 nM sensorgrams provided robust signal intensity while avoiding saturation observed at the highest analyte concentrations and were therefore used for consistent comparison among constructs.

### Functional cell-based assays reveal differential modulation of BMP-2 activity

To evaluate whether engineered BMP-2 binders modulate signaling in a cellular context, the alkaline phosphatase (ALP) activity was measured in C2C12 murine skeletal myoblasts, a well-established model of BMP-2–induced osteogenic differentiation. ALP activity was normalized to double-stranded DNA (dsDNA) content to account for differences in cell number across conditions (Fig. 6).

**Figure 6.**
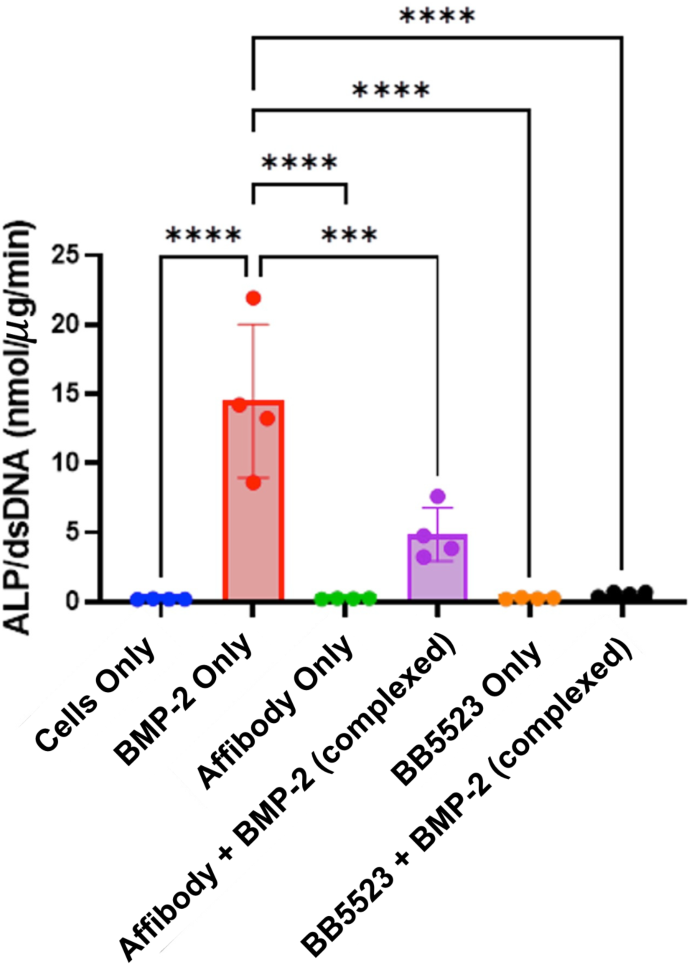
Cell-based alkaline phosphatase assays reveal differential modulation of BMP-2 signaling by engineered binders. ALP activity was measured in C2C12 myoblasts and normalized to dsDNA content following treatment with BMP-2 alone, engineered BB5523 binders, Affibody constructs, or their respective combinations with BMP-2. BMP-2 alone induces a robust osteogenic response relative to unstimulated controls. BB5523 binders and Affibody constructs alone show minimal ALP activity, comparable to baseline. When combined with BMP-2, both binder classes significantly attenuate BMP-2–driven ALP induction compared to BMP-2 alone. Notably, BB5523 produces a stronger reduction in BMP-2–mediated osteogenic differentiation than Affibody constructs under the conditions tested. Statistical significance was determined by one-way ANOVA (***p < 0.001, ****p < 0.0001).

Treatment with BMP-2 alone (50 ng mL ¹) induced a robust osteogenic response, with ALP activity markedly elevated relative to unstimulated cells. In contrast, treatment with either BB5523 binders or a BMP-2-specific affibody^11^ alone resulted in ALP levels comparable to baseline controls, indicating that neither binder class independently activates osteogenic signaling in this assay.

When combined with BMP-2, both BB5523 binders and the BMP-2-specific affibody significantly altered BMP-2–driven ALP induction relative to BMP-2 alone. Notably, ALP activity in all binder + BMP-2 conditions remained significantly lower than that observed with BMP-2 treatment alone, demonstrating that both binder classes attenuate BMP-2–mediated osteogenic differentiation under the tested conditions. Comparison between binder classes revealed that BB5523 more strongly suppressed BMP-2–induced ALP activity than Affibody constructs, indicating more effective inhibition of BMP-2 signaling in this cellular assay.

These functional differences are consistent with the intentional design of BB5523 binders within a lower-nanomolar affinity regime, which enables reversible and tunable modulation of BMP-2 activity rather than maximal growth factor signaling. In contrast, affibody constructs—previously reported to bind BMP-2 with high affinity—exhibited weaker attenuation of BMP-2–mediated ALP induction, consistent with partial preservation of BMP-2 bioactivity and previous results^34^ Together, these results demonstrate that BB5523 binders more effectively limit BMP-2–driven osteogenic differentiation in cells, highlighting their potential utility for precisely modulating BMP-2 signaling.

## Discussion

This study demonstrates that de novo protein design, when integrated with deep-learning refinement and experimental validation, can yield high-affinity binders targeting structurally complex growth factor epitopes^11^. Three key advances emerge from this work. First, we establish an integrated computational–experimental framework capable of addressing the challenges associated with designing binders to β-sheet–rich, receptor-engaging epitopes such as the BMP-2 knuckle region^12^. On one hand, the failure of Phase I designs to bind BMP-2 experimentally, despite favorable in silico metrics, highlights limitations of energy-based modeling for this class of interface^2413^. In this context, docking metrics were not predictive of experimental binding. On the other hand, the success of Phase II designs highlights the value of deep-learning–driven backbone refinement in correcting subtle geometric inaccuracies and improving interface complementarity beyond what physics-based approaches alone can capture^15^

Second, we demonstrate improved predictive reliability of the integrated design pipeline^15,16^ Computational assessments of structural integrity and interface geometry were generally consistent with experimental outcomes, including proper folding, high-affinity binding measured by bio-layer interferometry, and consistent functional responses in cell-based assays. The sharp contrast between Phase I and Phase II performance emphasizes the necessity of combining physics-based modeling with deep-learning refinement to achieve experimentally robust binders for challenging targets.

A central challenge underlying these results is the intrinsic bias of current deep–learning–based protein design models toward α-helical secondary structures^15^ Many diffusion- and sequence-based models are trained predominantly on helical folds, making accurate generation of extended β-sheet interfaces more difficult. This limitation is particularly pronounced for targets such as the BMP-2 knuckle epitope, which requires precise strand registry and hydrogen-bonding networks for effective binding^12^ In this context, partial diffusion^15^ and hybrid physics–deep-learning refinement^16,27^ were essentia for preserving β-sheet geometry while enabling localized backbone correction, helping to overcome a key structural limitation of current generative models.

Third, the engineered binders exhibit apparent binding affinities in the low-nanomolar regime. Notably, BB5523 exhibited an apparent equilibrium dissociation constant of 2.07 nM despite being discovered through a computational design campaign targeting a structurally challenging β-sheet-mediated interface. Despite achieving tight binding, the interaction remains tunable, as demonstrated by alanine substitutions that systematically modulate binding affinity^35^. Functional assays suggest that differences in biochemical binding strength translate into measurable differences in cellular signaling output, supporting a tunable relationship between molecular affinity and biological response. Effective control, therefore, arises not from maximal affinity alone, but from balancing specificity, reversibility, and binding strength within a defined therapeutic window.

Mutational analysis reveals that binding is partially governed by a dominant hotspot residue. In particular, substitution of T42 results in a substantial loss of binding affinity, indicating that this residue plays a central role in the interaction interface. In contrast, mutations at positions I44 and L46 produce more modest reductions in affinity, suggesting that these residues contribute to binding but do not dominate the interaction energetics. The retention of measurable binding in the triple mutant further indicates that the interaction is not solely driven by a small number of side-chain contacts, but instead involves a distributed interface with contributions from additional interactions, potentially including backbone-mediated complementarity or alternative binding modes not fully captured by the design model.

Notably, statistical comparison of kinetic fits across concentrations (P < 0.0001 for all constructs) indicates that a single global binding model does not fully describe the data. This suggests deviations from ideal 1:1 binding behavior in the BLI assay, highlighting the importance of cautious interpretation of fitted kinetic parameters.

This deviation from ideal 1:1 behavior suggests that BMP-2 engagement by the designed binders is influenced by non-ideal factors, including surface immobilization effects inherent to BLI assays, conformational flexibility at the interface, or heterogeneous binding modes^36^ Accordingly, kinetic parameters derived from global fitting are not interpreted as absolute thermodynamic constants. Instead, apparent K_D_ values derived from single-concentration fits are used as a consistent comparative framework to assess relative mutational effects. Because BMP-2 exists as a homodimer, multivalent engagement cannot be excluded; however, the present data do not permit unambiguous determination of binding stoichiometry^36^

Beyond validating the present designs, this framework provides a generalizable blueprint for next-generation binder development; however, several limitations and opportunities warrant consideration. The current binders reflect limitations in existing computational energy functions^23^, particularly in modeling extended β-sheet interactions, long-range electrostatics, conformational heterogeneity, and solvent-mediated effects at protein–protein interfaces. Observed differences in binding affinity compared to previously reported values highlight the sensitivity of BLI measurements to experimental conditions, including protein preparation, immobilization strategy, and concentration accuracy. These limitations likely contribute to discrepancies between predicted and experimentally observed binding behavior.

Future efforts may address these limitations through integration of experimental evolution and high-throughput mutagenesis to explore sequence space beyond what is readily accessible by design alone^37^. Such approaches would enable systematic optimization of affinity and stability while generating target-specific datasets to inform retraining or fine-tuning of data-driven models. Incorporation of ensemble-based design strategies, explicit modeling of alternative binding poses, electrostatics-aware solvation models, and improved treatment of β-strand register shifts may further reduce current sources of predictive uncertainty. Recent advances in AI-driven protein design workflows further highlight opportunities for improving binder discovery beyond the approaches used here^38^. Integrated design frameworks such as BindCraft combine diffusion-based backbone generation, sequence optimization, structural prediction and iterative filtering into scalable pipelines capable of producing experimentally validated binders across diverse targets^38^ Similarly, emerging all-atom deep-learning frameworks such as RoseTTAFold All-Atom have begun to bridge the gap between physics-based modeling and data-driven structure prediction by jointly modeling proteins, ligands and macromolecular interfaces^39^ Incorporation of these next-generation approaches may improve interface sampling, reduce false-positives, and provide more accurate ranking of candidate binders prior to experimental testing. Together, these developments suggest that future design campaigns will increasingly rely on hybrid workflows integrating generative AI methods, structure prediction and experimental feedback to improve success rates for challenging interfaces^28,38,39^

From a translational perspective, the next logical step is the co-design of binders and delivery systems. Incorporation of BMP-2 binders into biomaterial platforms, such as hydrogels^11,12^ nanoscaffolds, or injectable depots, could enable sustained, localized modulation of growth factor presentation while further decoupling therapeutic efficacy from bulk dosing and improving BMP-2 safety by lowering supraphysiological BMP-2 doses. Finally, in vivo validation will be essential to assess pharmacokinetics, immunogenicity, and long-term efficacy, and to determine whether computationally tuned affinity and reversibility translate into improved therapeutic windows under physiological conditions.

## Conclusion

This work demonstrates that the synergy of modern computational protein design, deep-learning refinement, and experimental validation can yield de novo protein binders with clinically relevant properties. The engineered BMP-2 binders combine high specificity with tunable affinity, with BB5523 achieving an apparent K_D_ of 2.07 nM, offering a protein-level strategy to modulate growth factor activity without reliance on high-dose therapies.

Beyond advancing a BMP-2–targeted solution, this study establishes a generalizable blueprint for precision-engineered protein delivery systems. Although demonstrated here using the BMP-2 knuckle epitope, this two-phase computational–experimental strategy is broadly applicable to other challenging growth factors and protein–protein interfaces characterized by complex β-sheet-mediated recognition. Together, these findings highlight the potential of computationally guided binder design in regenerative medicine and lay the groundwork for next-generation protein therapeutics that balance efficacy, safety, and control.

## Methods

### Structure Preparation and Relaxation

#### Structure retrieval and preprocessing

The crystal structure of human bone morphogenetic protein-2 (BMP-2) in complex with repulsive guidance molecule C (RGMC) was obtained from the Protein Data Bank (PDB ID: 4UI1)^40,41^. Prior to computational design, the structure was preprocessed using PyMOL. All non-essential components, including the RGMC binding partner, crystallographic water molecules, and small-molecule ligands, were removed to generate an isolated BMP-2 dimer suitable for subsequent modeling and binder design.

#### Rosetta-based structural relaxation

The preprocessed BMP-2 structure was subjected to the Rosetta Relax protocol implemented in PyRosetta^42–45^ During this step, side-chain rotamers and backbone torsions were sampled within the Rosetta FastRelax energy-minimization framework to relieve steric clashes and optimize packing interactions^24,45^ Hydrogen atoms were added to ensure chemical completeness. The relaxation procedure generated an ensemble of energetically minimized conformations, from which the lowest-energy structure was selected based on Rosetta scoring and used as the reference model for all downstream binder design.

### Binder Candidate Generation

#### Motif grafting and scaffold selection

Binder candidates were generated computationally using a β-strand motif grafting strategy, in which short β-strand motifs were transplanted onto structurally stable protein scaffolds^16,21^. Design efforts focused on the BMP-2 knuckle epitope (residues 192–196), a region critical for receptor engagement and well suited for β-sheet-mediated complementarity (Fig. 2). Scaffold backbones from Rocklin et al., Science (2017) were selected to support stable presentation of grafted motifs and formation of extended β-sheet interactions at the interface^46^

#### In silico prescreening

The initial design pool was subjected to rapid computational prescreening to eliminate designs with obvious structural flaws, such as severe steric clashes or nonphysical backbone geometries. Designs were retained only if they exhibited favorable Rosetta total energy scores computed using the ref2015 force field and maintained plausible β-strand geometry at the intended binding interface^47^. Rosetta total energy scores reflect a weighted combination of physics-based and statistical terms, including van der Waals interactions, hydrogen bonding, electrostatics, solvation, and backbone torsional preferences. These terms are parameterized using high-resolution experimental protein structures, such that lower total energy scores correspond to more physically realistic conformations.

### Docking and Interface Analysis

#### Global docking with RosettaDock

Surviving candidates were subjected to a two-stage RosettaDock protocol to sample and refine binding orientations relative to BMP-2. In the low-resolution centroid mode, binding poses were sampled extensively with side chains represented as simplified centroids^15^. Promising complexes were subsequently advanced to high-resolution, all-atom refinement, during which side-chain conformations and rigid-body orientations were optimized. Across the top-performing Phase I binders and the Phase II design BB5523, the lowest-RMSD models consistently remained below ~0.6 Å relative to the corresponding design model; accordingly, designs exhibiting backbone RMSD values within this range were retained for downstream evaluation.

### Structure Prediction and Validation

#### Deep-learning–based structure prediction

To assess fold fidelity and complex geometry independently of the Rosetta design process, monomeric structures were predicted using AlphaFold2, while binder-BMP-2 complex structures were evaluated using AlphaFold-Multimer^25,29^. These predictions provided orthogonal validation of structural integrity and binding orientation relative to the computational design models.

#### Evaluation criteria

Predicted structures were evaluated using multiple quantitative metrics. Designs were required to exhibit backbone RMSD values < ~3 Å relative to their corresponding design models, pLDDT confidence scores exceeding 80 across structured regions, and interface predicted aligned error (pAE) values below 10 Å. Candidates satisfying all criteria were advanced to deep-learning–driven refinement.

### Deep Learning–Driven Refinement

#### RFDiffusion optimization

Top-ranking Phase I designs were refined using partial RFDiffusion, a denoising diffusion–based framework for protein design^28^ Partial denoising enabled localized backbone reorganization while preserving the overall fold, thereby improving interface geometry, backbone stability, and packing interactions.

#### ProteinMPNN sequence design

Final sequence optimization was performed using the ProteinMPNN–FastRelax protocol^27,31^. Multiple sequence variants were sampled for each refined structural template, followed by Rosetta FastRelax to optimize side-chain packing. Designs were selected based on reductions in Rosetta total energy scores, improved interface complementarity, and favorable predicted structural properties.

### Protein purification

#### Transformation of Plasmids into E. coli

Plasmids encoding BB5523 and variant constructs were transformed into NEB Turbo competent E. coli cells following the manufacturer’s protocol. Briefly, competent cells were thawed on ice for 10 min, and 5 µL of plasmid DNA was added to the cell suspension. The mixture was gently mixed and incubated on ice for 30 min, followed by heat shock at 42 °C for 30 s. Cells were recovered in 950 µL of SOC medium and incubated at 37 °C with shaking for 60 min. Recovered cells were plated onto LB agar supplemented with kanamycin and incubated at 30 °C overnight.

#### Auto-Induction Expression of Protein Using AIM Medium

Plasmid sequences were verified by Sanger sequencing prior to expression. Glycerol stocks stored at −80 °C were streaked onto LB agar plates containing kanamycin and incubated overnight at 37 °C. A single colony was used to inoculate 5 mL LB supplemented with kanamycin and grown overnight at 37 °C with shaking. Starter cultures (100 µL) were transferred into 50 mL of auto-induction medium (AIM) supplemented with kanamycin in baffled flasks. Cultures were incubated at 37 °C until mid-log phase (OD ~0.6–0.8), then shifted to 30 °C and allowed to express protein overnight (~16 h) with shaking. Protein expression was induced automatically through metabolic depletion of glucose and subsequent lactose uptake, eliminating the need for manual IPTG induction.

#### Cell Lysis and Purification

Cells were harvested by centrifugation (10,000 rpm, 10 min), and pellets were stored at −20 °C prior to lysis. Pellets were resuspended in either 50 mM Tris-HCl (pH 8.0) or 50 mM HEPES (pH 6.0), each containing 100 mM NaCl, and lysed using BugBuster reagent (250 µL per 50 mL culture). Lysates were clarified by centrifugation (10,000 rpm, 20–30 min), and supernatants were applied to Ni-NTA resin for immobilized metal affinity chromatography (IMAC). Columns were equilibrated in a resuspension buffer, and bound proteins were eluted stepwise using imidazole concentrations of 25 mM, 50 mM, 200 mM, and 400 mM. Eluted fractions were analyzed by SDS-PAGE to assess purity and yield. Proteins were buffer-exchanged to PBS and stored.

### Site-Directed Mutagenesis

#### Generation of BB5523 variants

Single-point variants (T42A, T42N, I44A, L46A) and the triple mutant (T42A/I44A/L46A) were generated from the BB55223 expression plasmid using PCR-based mutagenesis. Complementary synthetic DNA oligonucleotides encoding the desired amino acid substitutions were designed with overlapping homology regions and assembled into the linearized expression vector using Gibson Assembly. Assembled plasmids were transformed into *E. coli*, and all constructs were confirmed by Sanger sequencing prior to protein expression.

### Biophysical Characterization

Purified proteins were buffer-exchanged into 50 mM Tris, 50 mM NaCl (pH 7.8) and characterized using complementary biophysical techniques. Molecular mass was confirmed by MALDI-TOF mass spectrometry, secondary structure and folding integrity were evaluated by CD spectroscopy, and binding kinetics and affinity toward BMP-2 were quantified by BLI using purified binders at 1 µM concentration.

For CD measurements, purified proteins were desalted into a 10 mM Tris buffer (pH ~7.4) using Zeba Spin Desalting Columns (7 kDa MWCO, Thermo Fisher Scientific) to remove NaCl prior to analysis. Triplicate spectra were collected for each sample over 190-250 nm at a scan read of 10 nm/s using a 1 mm path length cuvette, cumulated, and converted to mean residue ellipticity (MRE) for normalization. Protein concentrations were determined by averaged triplicate 280 nm reads on UV-Vis spectrophotometry. Protein concentrations used for MRE calculations are summarized in Table SI.1

### Functional Cell-Based Assays

#### C2C12 Culturing

C2C12 murine myoblasts were cultured in high-glucose Dulbecco’s modified Eagle’s medium (DMEM) supplemented with 10% fetal bovine serum (FBS), 10,000 µg/mL streptomycin, and 10,000 U/mL penicillin, per the manufacturer’s recommended protocols.

#### Alkaline Phosphatase Activity

C2C12 myoblasts were seeded in a 96-well plate at 20,000 cells/mL in DMEM with 10% FBS and streptomycin, and penicillin. The cells were allowed to adhere for 6 hours in the 10% FBS media. The media was replaced with DMEM supplemented with 1% FBS, streptomycin, and penicillin, and fortified with treatment groups (50 ng/mL BMP-2 (about 1.92 nM) only, 3.84 nM high-affinity BMP-2-specific affibody, 3.84 nM BB5523, 1.92 nM BMP-2 mixed with 3.84 nM high-affinity BMP-2-specific affibody, or 1.92 nM BMP-2 mixed with 3.84 nM BB5523). The cells were incubated at 37 °C for 72 hours. After 72 hours, cells were lysed, and their ALP activity was quantified using a colorimetric assay, as described previously^11^ Briefly, a solution consisting of 2-amino-2methyl-1-propanol, p-nitrophenyl phosphate, and MgCl2 hexahydrate was mixed with the cell lysate for 20 minutes in the dark, and measuring the absorbance at 405 nm. The results were normalized to double stranded DNA (dsDNA) content for each well using the QuantiFluor dsDNA kit (Promega).

## Acknowledgments

This work was supported by grants from the NSF 2137880 to PH (total of $585,000) and NIH (R35-GM147507 MIRA to MHH). We gratefully acknowledge the University of Oregon Knight Campus facilities for providing technical resources and instrumentation essential to this project. We also thank our collaborators and trainees for their invaluable contributions, discussions, and support throughout the course of this study.

## Data Storage

All the computational assays are in https://github.com/ParisaH-Lab/publications/tree/main/BMP2_binders

All computational workflows were executed using standardized, version-controlled pipelines deployed on high-performance computing clusters. This ensured reproducibility across independent runs and enabled efficient iteration as new design insights were incorporated.

The raw data are provided as SI: CD and BLI.

## Notes

https://github.com/ParisaH-Lab/publications/tree/main/BMP2_binders

### Competing Interest Statement

The authors have declared no competing interest.

## References

(1) Wu, A.-M.; Bisignano, C.; James, S. L.; Abady, G. G.; Abedi, A.; Abu-Gharbieh, E.; Alhassan, R. K.; Alipour, V.; Arabloo, J.; Asaad, M.; Asmare, W. N.; Awedew, A. F.; Banach, M.; Banerjee, S. K.; Bijani, A.; Birhanu, T. T. M.; Bolla, S. R.; Cámera, L. A.; Chang, J.-C.; Cho, D. Y.; Chung, M. T.; Couto, R. A. S.; Dai, X.; Dandona, L.; Dandona, R.; Farzadfar, F.; Filip, I.; Fischer, F.; Fomenkov, A. A.; Gill, T. K.; Gupta, B.; Haagsma, J. A.; Haj-Mirzaian, A.; Hamidi, S.; Hay, S. I.; Ilic, I. M.; Ilic, M. D.; Ivers, R. Q.; Jürisson, M.; Kalhor, R.; Kanchan, T.; Kavetskyy, T.; Khalilov, R.; Khan, E. A.; Khan, M.; Kneib, C. J.; Krishnamoorthy, V.; Kumar, G. A.; Kumar, N.; Lalloo, R.; Lasrado, S.; Lim, S. S.; Liu, Z.; Manafi, A.; Manafi, N.; Menezes, R. G.; Meretoja, T. J.; Miazgowski, B.; Miller, T. R.; Mohammad, Y.; Mohammadian-Hafshejani, A.; Mokdad, A. H.; Murray, C. J. L.; Naderi, M.; Naimzada, M. D.; Nayak, V. C.; Nguyen, C. T.; Nikbakhsh, R.; Olagunju, A. T.; Otstavnov, N.; Otstavnov, S. S.; Padubidri, J. R.; Pereira, J.; Pham, H. Q.; Pinheiro, M.; Polinder, S.; Pourchamani, H.; Rabiee, N.; Radfar, A.; Rahman, M. H. U.; Rawaf, D. L.; Rawaf, S.; Saeb, M. R.; Samy, A. M.; Sanchez Riera, L.; Schwebel, D. C.; Shahabi, S.; Shaikh, M. A.; Soheili, A.; Tabarés-Seisdedos, R.; Tovani-Palone, M. R.; Tran, B. X.; Travillian, R. S.; Valdez, P. R.; Vasankari, T. J.; Velazquez, D. Z.; Venketasubramanian, N.; Vu, G. T.; Zhang, Z.-J.; Vos, T. Global, Regional, and National Burden of Bone Fractures in 204 Countries and Territories, 1990–2019: A Systematic Analysis from the Global Burden of Disease Study 2019. The Lancet Healthy Longevity 2021 2 (9), e580–e592. 10.1016/S2666-7568(21)00172-0.

(2) Thomas, J. D.; Kehoe, J. L. Bone Nonunion; StatPearls, 2023.

(3) Nicholson, J.; Makaram, N.; Simpson, A.; Keating, J. Fracture Nonunion in Long Bones: A Literature Review of Risk Factors and Surgical Management. Injury 2021 52, S3–S11. 10.1016/j.injury.2020.11.029.

(4) Schmidt, A. H. Autologous Bone Graft: Is It Still the Gold Standard? Injury 2021 52 S18–S22. 10.1016/j.injury.2021.01.043.

(5) Urist, M. R. Bone: Formation by Autoinduction. Science 1965 150 (3698), 893–899. 10.1126/science.150.3698.893.

(6) Halloran, D.; Durbano, H. W.; Nohe, A. Bone Morphogenetic Protein-2 in Development and Bone Homeostasis. JDB 2020 8 (3), 19. 10.3390/jdb8030019.

(7) Vantucci, C. E.; Krishan, L.; Cheng, A.; Prather, A.; Roy, K.; Guldberg, R. E. BMP-2 Delivery Strategy Modulates Local Bone Regeneration and Systemic Immune Responses to Complex Extremity Trauma. Biomater. Sci. 2021 9 (5), 1668–1682. 10.1039/D0BM01728K.

(8) Qi, J.; Wu, H.; Liu, G. Novel Strategies for Spatiotemporal and Controlled BMP-2 Delivery in Bone Tissue Engineering. Cell Transplant 2024 33, 09636897241276733. 10.1177/09636897241276733.

(9) Miyazono, K.; Kamiya, Y.; Morikawa, M. Bone Morphogenetic Protein Receptors and Signal Transduction. Journal of Biochemistry 2010 147 (1), 35–51. 10.1093/jb/mvp148.

(10) Zou, M.-L.; Chen, Z.-H.; Teng, Y.-Y.; Liu, S.-Y.; Jia, Y.; Zhang, K.-W.; Sun, Z.-L.; Wu, J.-J.; Yuan, Z.-D.; Feng, Y.; Li, X.; Xu, R.-S.; Yuan, F.-L. The Smad Dependent TGF-β and BMP Signaling Pathway in Bone Remodeling and Therapies. Front. Mol. Biosci. 2021 8 593310. 10.3389/fmolb.2021.593310.

(11) Dorogin, J.; Hochstatter, H. B.; Shepherd, S. O.; Svendsen, J. E.; Benz, M. A.; Powers, A. C.; Fear, K. M.; Townsend, J. M.; Prell, J. S.; Hosseinzadeh, P.; Hettiaratchi, M. H. Moderate-Affinity Affibodies Modulate the Delivery and Bioactivity of Bone Morphogenetic Protein-2. Adv Healthcare Materials 2023 12 (26), 2300793. 10.1002/adhm.202300793.

(12) Svendsen, J. E.; Ford, M. R.; Asnes, C. L.; Oh, S. C.; Dorogin, J.; Fear, K. M.; O’Hara-Smith, J. R.; Chisholm, L. O.; Phillips, S. R.; Harms, M. J.; Hosseinzadeh, P.; Hettiaratchi, M. H. Applying Computational Protein Design to Engineer Affibodies for Affinity-Controlled Delivery of Vascular Endothelial Growth Factor and Platelet-Derived Growth Factor. Biomacromolecules 2025 26 (6), 3463–3480. 10.1021/acs.biomac.5c00097.

(13) Zhao, G.; Zhang, L.; Che, L.; Li, H.; Liu, Y.; Fang, J. Revisiting Bone Morphogenetic Protein-2 Knuckle Epitope and Redesigning the Epitope-derived Peptides. Journal of Peptide Science 2021 27 (6), e3309. 10.1002/psc.3309.

(14) Sahtoe, D. D.; Coscia, A.; Mustafaoglu, N.; Miller, L. M.; Olal, D.; Vulovic, I.; Yu, T.-Y.; Goreshnik, I.; Lin, Y.-R.; Clark, L.; Busch, F.; Stewart, L.; Wysocki, V. H.; Ingber, D. E.; Abraham, J.; Baker, D. Transferrin Receptor Targeting by de Novo Sheet Extension. Proc. Natl. Acad. Sci. U.S.A 2021 118 (17), e2021569118. 10.1073/pnas.2021569118.

(15) Chaudhury, S.; Berrondo, M.; Weitzner, B. D.; Muthu, P.; Bergman, H.; Gray, J. J. Benchmarking and Analysis of Protein Docking Performance in Rosetta v3.2. PLoS ONE 2011 6 (8), e22477. 10.1371/journal.pone.0022477.

(16) Silva, D.-A.; Correia, B. E.; Procko, E. Motif-Driven Design of Protein-Protein Interfaces. Methods Mol. Biol 2016 1414, 285–304. 10.1007/978-1-4939-3569-7_17.

(17) Chaudhury, S.; Lyskov, S.; Gray, J. J. PyRosetta: A Script-Based Interface for Implementing Molecular Modeling Algorithms Using Rosetta. Bioinformatics 2010 26 (5), 689–691. 10.1093/bioinformatics/btq007.

(18) Leaver-Fay, A.; Tyka, M.; Lewis, S. M.; Lange, O. F.; Thompson, J.; Jacak, R.; Kaufman, K. W.; Renfrew, P. D.; Smith, C. A.; Sheffler, W.; Davis, I. W.; Cooper, S.; Treuille, A.; Mandell, D. J.; Richter, F.; Ban, Y.-E. A.; Fleishman, S. J.; Corn, J. E.; Kim, D. E.; Lyskov, S.; Berrondo, M.; Mentzer, S.; Popović, Z.; Havranek, J. J.; Karanicolas, J.; Das, R.; Meiler, J.; Kortemme, T.; Gray, J. J.; Kuhlman, B.; Baker, D.; Bradley, P. Rosetta3. In Methods in Enzymology; Elsevier, 2011; Vol. 487, pp 545–574. 10.1016/B978-0-12-381270-4.00019-6.

(19) Cock, P. J. A.; Antao, T.; Chang, J. T.; Chapman, B. A.; Cox, C. J.; Dalke, A.; Friedberg, I.; Hamelryck, T.; Kauff, F.; Wilczynski, B.; De Hoon, M. J. L. Biopython: Freely Available Python Tools for Computational Molecular Biology and Bioinformatics. Bioinformatics 2009 25 (11), 1422–1423. 10.1093/bioinformatics/btp163.

(20) Hamelryck, T.; Manderick, B. PDB File Parser and Structure Class Implemented in Python. Bioinformatics 2003 19 (17), 2308–2310. 10.1093/bioinformatics/btg299.

(21) Fleishman, S. J.; Leaver-Fay, A.; Corn, J. E.; Strauch, E.-M.; Khare, S. D.; Koga, N.; Ashworth, J.; Murphy, P.; Richter, F.; Lemmon, G.; Meiler, J.; Baker, D. RosettaScripts: A Scripting Language Interface to the Rosetta Macromolecular Modeling Suite. PLoS ONE 2011 6 (6), e20161. 10.1371/journal.pone.0020161.

(22) Jones, D. T. Protein Secondary Structure Prediction Based on Position-Specific Scoring Matrices 1 1Edited by G. Von Heijne. Journal of Molecular Biology 1999 292 (2), 195–202. 10.1006/jmbi.1999.3091.

(23) Maguire, J. B.; Haddox, H. K.; Strickland, D.; Halabiya, S. F.; Coventry, B.; Griffin, J. R.; Pulavarti, S. V. S. R. K.; Cummins, M.; Thieker, D. F.; Klavins, E.; Szyperski, T.; DiMaio, F.; Baker, D.; Kuhlman, B. Perturbing the Energy Landscape for Improved Packing during Computational Protein Design. Proteins 2021 89 (4), 436–449. 10.1002/prot.26030.

(24) Alford, R. F.; Leaver-Fay, A.; Jeliazkov, J. R.; O’Meara, M. J.; DiMaio, F. P.; Park, H.; Shapovalov, M. V.; Renfrew, P. D.; Mulligan, V. K.; Kappel, K.; Labonte, J. W.; Pacella, M. S.; Bonneau, R.; Bradley, P.; Dunbrack, R. L.; Das, R.; Baker, D.; Kuhlman, B.; Kortemme, T.; Gray, J. J. The Rosetta All-Atom Energy Function for Macromolecular Modeling and Design. J. Chem. Theory Comput. 2017 13 (6), 3031–3048. 10.1021/acs.jctc.7b00125.

(25) Evans, R.; O’Neill, M.; Pritzel, A.; Antropova, N.; Senior, A.; Green, T.; Žídek, A.; Bates, R.; Blackwell, S.; Yim, J.; Ronneberger, O.; Bodenstein, S.; Zielinski, M.; Bridgland, A.; Potapenko, A.; Cowie, A.; Tunyasuvunakool, K.; Jain, R.; Clancy, E.; Kohli, P.; Jumper, J.; Hassabis, D. Protein Complex Prediction with AlphaFold-Multimer. Bioinformatics October 4, 2021. 10.1101/2021.10.04.463034.

(26) Chennamsetty, N.; Voynov, V.; Kayser, V.; Helk, B.; Trout, B. L. Design of Therapeutic Proteins with Enhanced Stability. Proc. Natl. Acad. Sci. U.S.A. 2009 106 (29), 11937–11942. 10.1073/pnas.0904191106.

(27) Dauparas, J.; Anishchenko, I.; Bennett, N.; Bai, H.; Ragotte, R. J.; Milles, L. F.; Wicky, B. I. M.; Courbet, A.; De Haas, R. J.; Bethel, N.; Leung, P. J. Y.; Huddy, T. F.; Pellock, S.; Tischer, D.; Chan, F.; Koepnick, B.; Nguyen, H.; Kang, A.; Sankaran, B.; Bera, A. K.; King, N. P.; Baker, D. Robust Deep Learning–Based Protein Sequence Design Using ProteinMPNN. Science 2022 378 (6615), 49–56. 10.1126/science.add2187.

(28) Watson, J. L.; Juergens, D.; Bennett, N. R.; Trippe, B. L.; Yim, J.; Eisenach, H. E.; Ahern, W.; Borst, A. J.; Ragotte, R. J.; Milles, L. F.; Wicky, B. I. M.; Hanikel, N.; Pellock, S. J.; Courbet, A.; Sheffler, W.; Wang, J.; Venkatesh, P.; Sappington, I.; Torres, S. V.; Lauko, A.; De Bortoli, V.; Mathieu, E.; Ovchinnikov, S.; Barzilay, R.; Jaakkola, T. S.; DiMaio, F.; Baek, M.; Baker, D. De Novo Design of Protein Structure and Function with RFdiffusion. Nature 2023 620 (7976), 1089–1100. 10.1038/s41586-023-06415-8.

(29) Jumper, J.; Evans, R.; Pritzel, A.; Green, T.; Figurnov, M.; Ronneberger, O.; Tunyasuvunakool, K.; Bates, R.; Žídek, A.; Potapenko, A.; Bridgland, A.; Meyer, C.; Kohl, S. A. A.; Ballard, A. J.; Cowie, A.; Romera-Paredes, B.; Nikolov, S.; Jain, R.; Adler, J.; Back, T.; Petersen, S.; Reiman, D.; Clancy, E.; Zielinski, M.; Steinegger, M.; Pacholska, M.; Berghammer, T.; Bodenstein, S.; Silver, D.; Vinyals, O.; Senior, A. W.; Kavukcuoglu, K.; Kohli, P.; Hassabis, D. Highly Accurate Protein Structure Prediction with AlphaFold. Nature 2021 596 (7873), 583–589. 10.1038/s41586-021-03819-2.

(30) Cao, L.; Coventry, B.; Goreshnik, I.; Huang, B.; Sheffler, W.; Park, J. S.; Jude, K. M.; Marković, I.; Kadam, R. U.; Verschueren, K. H. G.; Verstraete, K.; Walsh, S. T. R.; Bennett, N.; Phal, A.; Yang, A.; Kozodoy, L.; DeWitt, M.; Picton, L.; Miller, L.; Strauch, E.-M.; DeBouver, N. D.; Pires, A.; Bera, A. K.; Halabiya, S.; Hammerson, B.; Yang, W.; Bernard, S.; Stewart, L.; Wilson, I. A.; Ruohola-Baker, H.; Schlessinger, J.; Lee, S.; Savvides, S. N.; Garcia, K. C.; Baker, D. Design of Protein-Binding Proteins from the Target Structure Alone. Nature 2022 605 (7910), 551–560. 10.1038/s41586-022-04654-9.

(31) Bennett, N. R.; Coventry, B.; Goreshnik, I.; Huang, B.; Allen, A.; Vafeados, D.; Peng, Y. P.; Dauparas, J.; Baek, M.; Stewart, L.; DiMaio, F.; De Munck, S.; Savvides, S. N.; Baker, D. Improving de Novo Protein Binder Design with Deep Learning. Nat Commun 2023 14 (1), 2625. 10.1038/s41467-023-38328-5.

(32) Micsonai, A.; Wien, F.; Murvai, N.; Nyiri, M. P.; Balatoni, B.; Lee, Y.-H.; Molnár, T.; Goto, Y.; Jamme, F.; Kardos, J. BeStSel: Analysis Site for Protein CD Spectra—2025 Update. Nucleic Acids Research 2025 53 (W1), W73–W83. 10.1093/nar/gkaf378.

(33) Micsonai, A.; Wien, F.; Kernya, L.; Lee, Y.-H.; Goto, Y.; Réfrégiers, M.; Kardos, J. Accurate Secondary Structure Prediction and Fold Recognition for Circular Dichroism Spectroscopy. Proc. Natl. Acad. Sci. U.S.A. 2015 112 (24). 10.1073/pnas.1500851112.

(34) Dorogin, J.; Hochstatter, H. B.; Shepherd, S. O.; Svendsen, J. E.; Benz, M. A.; Powers, A. C.; Fear, K. M.; Townsend, J. M.; Prell, J. S.; Hosseinzadeh, P.; Hettiaratchi, M. H. Moderate-Affinity Affibodies Modulate the Delivery and Bioactivity of Bone Morphogenetic Protein-2. Adv Healthcare Materials 2023 12 (26), 2300793. 10.1002/adhm.202300793.

(35) Hettiaratchi, M. H.; Krishnan, L.; Rouse, T.; Chou, C.; McDevitt, T. C.; Guldberg, R. E. Heparin-Mediated Delivery of Bone Morphogenetic Protein-2 Improves Spatial Localization of Bone Regeneration. Sci. Adv 2020 6 (1), eaay1240. 10.1126/sciadv.aay1240.

(36) Myszka, D. G. Improving Biosensor Analysis. J Mol. Recognit. 1999 12 (5), 279–284. 10.1002/(SICI)1099-1352(199909/10)12:5%3C279::AID-JMR473%3E3.0.CO;2-3.

(37) McConnell, A.; Hackel, B. J. Protein Engineering via Sequence-Performance Mapping. Cell Systems 2023 14 (8), 656–666. 10.1016/j.cels.2023.06.009.

(38) Pacesa, M.; Nickel, L.; Schellhaas, C.; Schmidt, J.; Pyatova, E.; Kissling, L.; Barendse, P.; Choudhury, J.; Kapoor, S.; Alcaraz-Serna, A.; Cho, Y.; Ghamary, K. H.; Vinué, L.; Yachnin, B. J.; Wollacott, A. M.; Buckley, S.; Westphal, A. H.; Lindhoud, S.; Georgeon, S.; Goverde, C. A.; Hatzopoulos, G. N.; Gönczy, P.; Muller, Y. D.; Schwank, G.; Swarts, D. C.; Vecchio, A. J.; Schneider, B. L.; Ovchinnikov, S.; Correia, B. E. One-Shot Design of Functional Protein Binders with BindCraft Nature 2025 646 (8084), 483–492. 10.1038/s41586-025-09429-6.

(39) Krishna, R.; Wang, J.; Ahern, W.; Sturmfels, P.; Venkatesh, P.; Kalvet, I.; Lee, G. R.; Morey-Burrows, F. S.; Anishchenko, I.; Humphreys, I. R.; McHugh, R.; Vafeados, D.; Li, X.; Sutherland, G. A.; Hitchcock, A.; Hunter, C. N.; Kang, A.; Brackenbrough, E.; Bera, A. K.; Baek, M.; DiMaio, F.; Baker, D. Generalized Biomolecular Modeling and Design with RoseTTAFold All-Atom. Science 2024 384 (6693), eadl2528. 10.1126/science.adl2528.

(40) Berman, H. M. The Protein Data Bank. Nucleic Acids Research 2000 28 (1), 235–242. 10.1093/nar/28.1.235.

(41) Healey, E. G.; Bishop, B.; Elegheert, J.; Bell, C. H.; Padilla-Parra, S.; Siebold, C. Repulsive Guidance Molecule Is a Structural Bridge between Neogenin and Bone Morphogenetic Protein. Nat Struct Mol Biol 2015 22 (6), 458–465. 10.1038/nsmb.3016.

(42) Tyka, M. D.; Keedy, D. A.; André, I.; DiMaio, F.; Song, Y.; Richardson, D. C.; Richardson, J. S.; Baker, D. Alternate States of Proteins Revealed by Detailed Energy Landscape Mapping. Journal of Molecular Biology 2011 405 (2), 607–618. 10.1016/j.jmb.2010.11.008.

(43) Khatib, F.; Cooper, S.; Tyka, M. D.; Xu, K.; Makedon, I.; Popovic, Z.; Baker, D.; Players, F. Algorithm Discovery by Protein Folding Game Players. Proc Natl Acad Sci U S A 2011 108 (47), 18949–18953. 10.1073/pnas.1115898108.

(44) Nivón, L. G.; Moretti, R.; Baker, D. A Pareto-Optimal Refinement Method for Protein Design Scaffolds. PLoS ONE 2013 8 (4), e59004. 10.1371/journal.pone.0059004.

(45) Conway, P.; Tyka, M. D.; DiMaio, F.; Konerding, D. E.; Baker, D. Relaxation of Backbone Bond Geometry Improves Protein Energy Landscape Modeling. Protein Science 2014 23 (1), 47–55. 10.1002/pro.2389.

(46) Rocklin, G. J.; Chidyausiku, T. M.; Goreshnik, I.; Ford, A.; Houliston, S.; Lemak, A.; Carter, L.; Ravichandran, R.; Mulligan, V. K.; Chevalier, A.; Arrowsmith, C. H.; Baker, D. Global Analysis of Protein Folding Using Massively Parallel Design, Synthesis, and Testing. Science 2017 357 (6347), 168–175. 10.1126/science.aan0693.

(47) Leman, J. K.; Weitzner, B. D.; Lewis, S. M.; Adolf-Bryfogle, J.; Alam, N.; Alford, R. F.; Aprahamian, M.; Baker, D.; Barlow, K. A.; Barth, P.; Basanta, B.; Bender, B. J.; Blacklock, K.; Bonet, J.; Boyken, S. E.; Bradley, P.; Bystroff, C.; Conway, P.; Cooper, S.; Correia, B. E.; Coventry, B.; Das, R.; De Jong, R. M.; DiMaio, F.; Dsilva, L.; Dunbrack, R.; Ford, A. S.; Frenz, B.; Fu, D. Y.; Geniesse, C.; Goldschmidt, L.; Gowthaman, R.; Gray, J. J.; Gront, D.; Guffy, S.; Horowitz, S.; Huang, P.-S.; Huber, T.; Jacobs, T. M.; Jeliazkov, J. R.; Johnson, D. K.; Kappel, K.; Karanicolas, J.; Khakzad, H.; Khar, K. R.; Khare, S. D.; Khatib, F.; Khramushin, A.; King, I. C.; Kleffner, R.; Koepnick, B.; Kortemme, T.; Kuenze, G.; Kuhlman, B.; Kuroda, D.; Labonte, J. W.; Lai, J. K.; Lapidoth, G.; Leaver-Fay, A.; Lindert, S.; Linsky, T.; London, N.; Lubin, J. H.; Lyskov, S.; Maguire, J.; Malmström, L.; Marcos, E.; Marcu, O.; Marze, N. A.; Meiler, J.; Moretti, R.; Mulligan, V. K.; Nerli, S.; Norn, C.; Ó’Conchúir, S.; Ollikainen, N.; Ovchinnikov, S.; Pacella, M. S.; Pan, X.; Park, H.; Pavlovicz, R. E.; Pethe, M.; Pierce, B. G.; Pilla, K. B.; Raveh, B.; Renfrew, P. D.; Burman, S. S. R.; Rubenstein, A.; Sauer, M. F.; Scheck, A.; Schief, W.; Schueler-Furman, O.; Sedan, Y.; Sevy, A. M.; Sgourakis, N. G.; Shi, L.; Siegel, J. B.; Silva, D.-A.; Smith, S.; Song, Y.; Stein, A.; Szegedy, M.; Teets, F. D.; Thyme, S. B.; Wang, R. Y.-R.; Watkins, A.; Zimmerman, L.; Bonneau, R. Macromolecular Modeling and Design in Rosetta: Recent Methods and Frameworks. Nat Methods 2020 17 (7), 665–680. 10.1038/s41592-020-0848-2.

