## Supporting Information for "From Code to Cure: Computationally Designed BMP-2 Binders Using AI-Integrated Pipelines for Controlled Bone Regeneration"

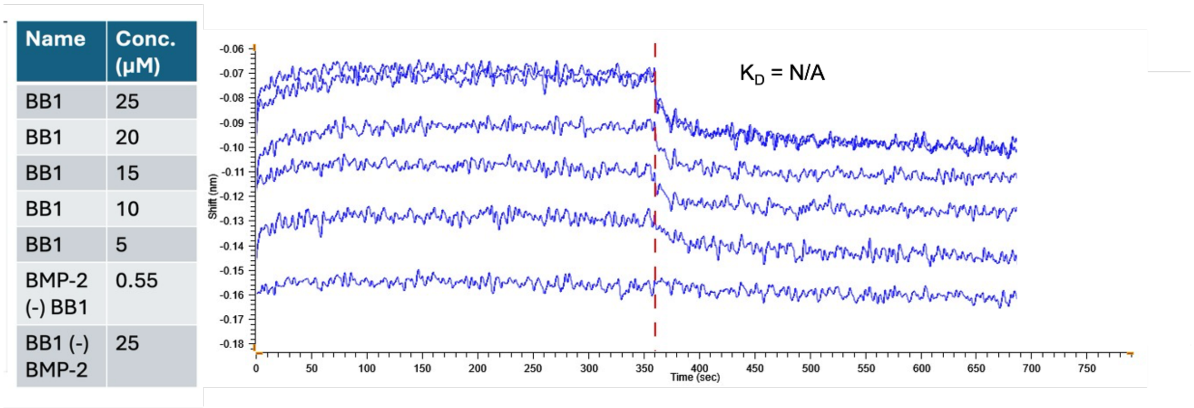

| Name | Conc. (μM) |
|---|---|
| BB1 | 25 |
| BB1 | 20 |
| BB1 | 15 |
| BB1 | 10 |
| BB1 | 5 |
| BMP-2 (-) BB1 | 0.55 |
| BB1 (-) BMP-2 | 25 |



**Figure SI.1 | BLI of Phase I binder shows inconclusive results of binding over negative control.** Representative BLI sensorgrams showing association and dissociation kinetics of Phase I design BB1 binding to BMP-2. The association phase (0–600 s) is followed by dissociation upon transfer to the buffer (vertical dashed line). Global fitting of the sensorgrams yielded no conclusive $K_D$ values, indicating no reproducible binding. Together, these data informed the refinement of the workflow to produce stronger binders.

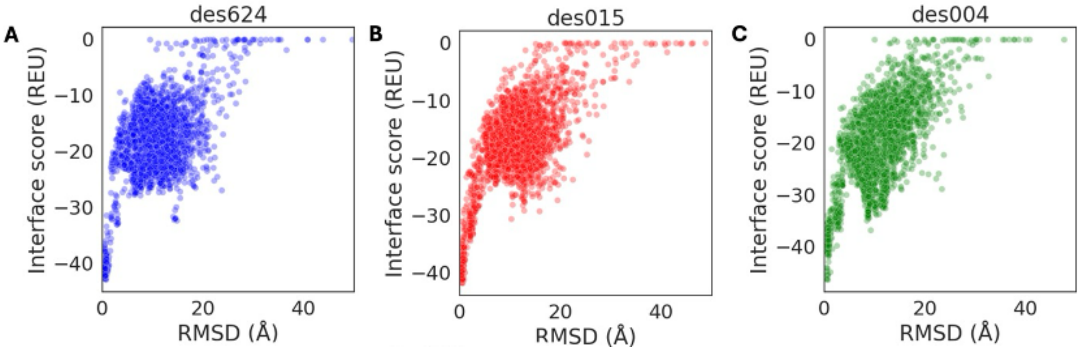



**Figure SI.2 | Additional RosettaDock energy landscapes for Phase I BMP-2 binder candidates.** Docking simulations for Phase I binders des624, des015, and des004 against BMP-2. Each point represents a docked decoy complex generated by RosettaDock and is plotted as interface energy score (Isc) versus RMSD relative to the designed binding pose. Several candidates exhibit convergent binding funnels characterized by lower interface energies at reduced RMSD values. These data complement the representative docking examples shown in Figure 3 and illustrate the range of docking behaviors observed among Phase I designs.

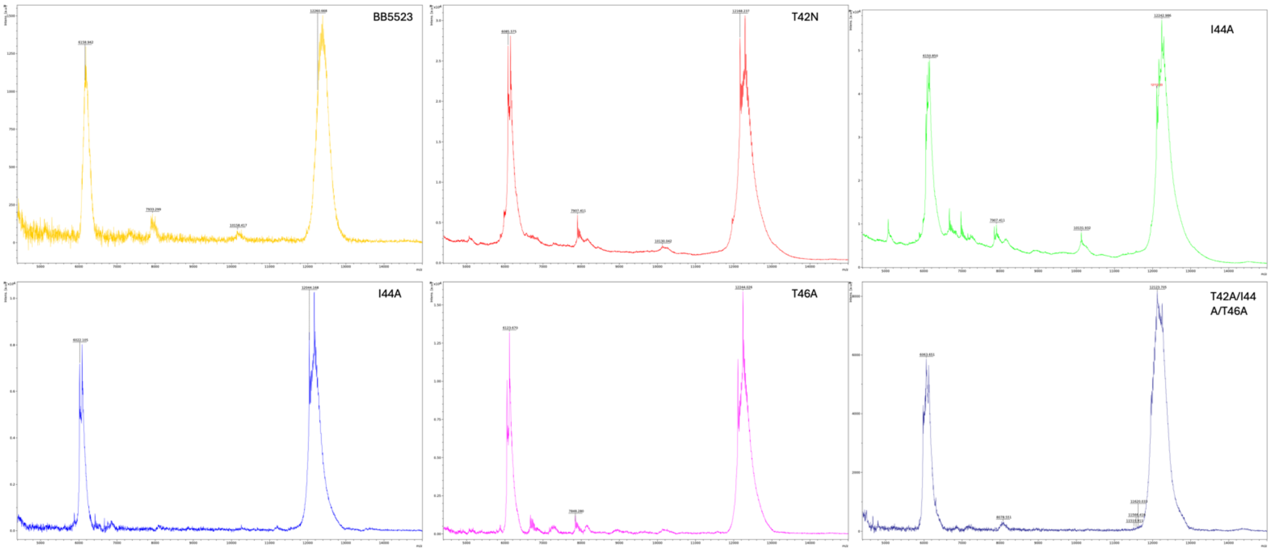


**Figure SI.3 | MALDI-TOF mass spectrometry confirms molecular integrity of designed BMP-2 binders.** Measured molecular weights of BB5523 and five mutated binder variants closely match predicted values (~12.3 kDa), demonstrating correct expression, sequence fidelity, and structural integrity of the recombinant proteins.

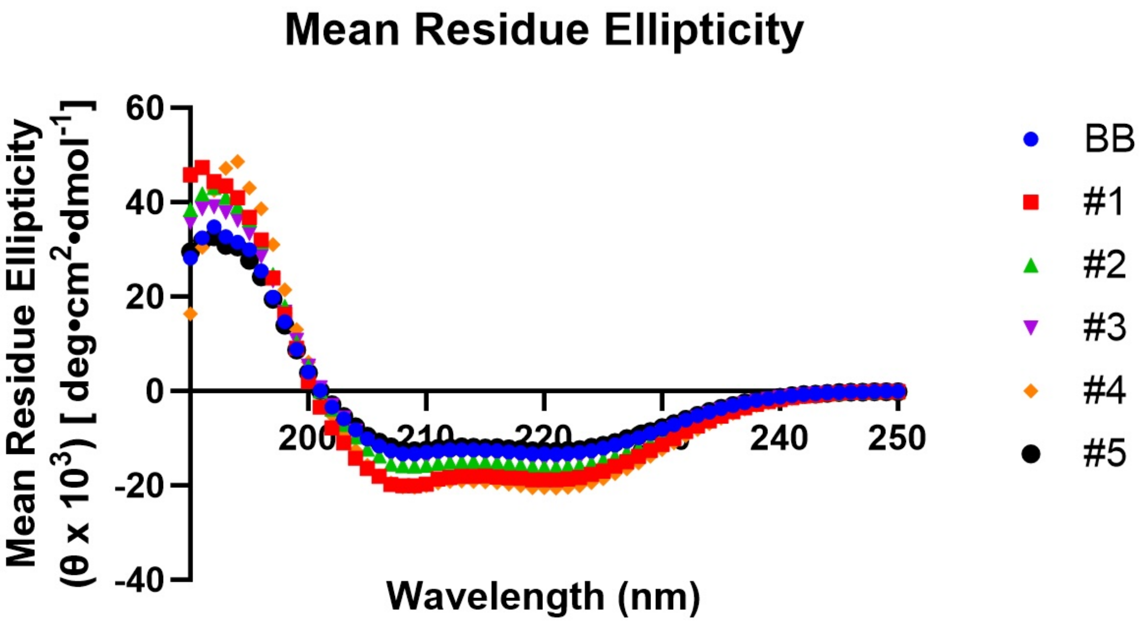


**Figure SI.4 | Circular dichroism spectroscopy confirms predominantly α-helical secondary structure of BB5523 and variants.** Far-UV CD spectra (190–250 nm) display characteristic minima at ~208 and ~222 nm and highly consistent overlays across independently measured samples, indicating stable folding and reproducible secondary structure across constructs. Quantitative secondary-structure estimates are provided in Table SI.2 BB = BB5523, #1 = T42A, #2 = T42N, #3 = I44A, #4 = L46A, #5 = T42A/I44A/L46A.

| Name | MWs (Da) | Conc (g/L) | Conc (M) |
|---|---|---|---|
| BB | 12265.65 | 0.348 | 2.83719E-05 |
| T42A | 11560.78 | 0.134 | 1.15909E-05 |
| T42N | 12169.27 | 0.272 | 2.23514E-05 |
| I44A | 12114.4 | 0.263 | 2.17097E-05 |
| L46A | 12250.88 | 0.258 | 2.10597E-05 |
| T42A/I44A/L46A | 12045.11 | 0.269 | 2.23327E-05 |

**Table SI.1 | Molecular weights and concentrations of BB5523 and variant constructs used for circular dichroism (CD) measurements.** Protein concentrations were calculated from measured mass concentrations and theoretical molecular weights and were used for conversion of CD spectra to mean residue ellipticity (MRE).

| Construct | Helix (%) | Antiparallel (%) | Parallel (%) | Turn (%) | Others (%) | NRMSD |
|---|---|---|---|---|---|---|
| **BB5523** | **28.4** | **14.8** | **5.1** | **12.2** | **39.4** | **0.00584** |
| **T42A** | **47.0** | **6.5** | **0.0** | **11.7** | **34.8** | **0.00832** |
| **T42N** | **34.4** | **14.0** | **3.6** | **12.0** | **36.0** | **0.00627** |
| **I44A** | **30.4** | **15.4** | **4.1** | **12.4** | **37.7** | **0.00578** |
| **L46A** | **53.8** | **10.8** | **0.0** | **5.2** | **30.2** | **0.02494** |
| **T42A/I44A/L46A** | **26.5** | **15.0** | **6.8** | **11.9** | **39.7** | **0.00562** |

**Table SI.2 | BeStSel secondary structure estimates for BB5523 and variant constructs.**

### DNA sequences:

Des004:
CTCTAGAAATAATTTTGTTTAACTTTAAGAAGGAGATATACCATGGGCTCGATCGAAGAGAAA
GCTAAGAAAATTATAGAGAAACTTTGGCAGGAAGCTAAGGAAAAAGGAAAGAGCGAGGCTT
GGGCGGCGATGGAAGCAATTCAGCGCATCGTGCCGTGGGATTTAGCAGCAACCATCATAGT
TGAAGACGAAGAAATAGCTAAAAAAATAAAAGAAATTATTAAGAAAAAAGTGCCCGAGGCTTA

CATTGTGGTTACAGCAAACATAATAGTGATCAATGCACAAACGGAGGAATTATTAAAAATCGC
ATTAGAAGAAGCAAGAAAAATTTTCCTCGAGCACCACCACCACC

Des015:

CTCTAGAAATAATTTTGTTTAACTTTAAGAAGGAGATATACCATGGGCGATGAAGAGGAAAAG
AGAGCTCAAGAGATAGTCAAGAAACTTCGGGAAGAGTTAAAAAAGAAAGGTGCTTCAGAAG
AACAAATTCTTCGCTTGGCCGCTCAACAGATCCTTTGGCTTTTGAGAGCGGGTATAGTCTTT
ATTAGCTCAGATGAAAGAGAAGCGAGAAAAATTTCTGAAATTCTGAAGAAATTAGCACCTGA
GGTGCGCGTTGAAGTGAGAGATGGGATAGTTGTAGTACATGCACGGTCCGAAGAATTGAAG
AAAAAGCTGCATGAATTACTGAAGAAGGCAGCTCTCGAGCACCACCACCACC

Des004:
CTCTAGAAATAATTTTGTTTAACTTTAAGAAGGAGATATACCATGGGCTCGATCGAAGAGAAA
GCTAAGAAAATTATAGAGAAACTTTGGCAGGAAGCTAAGGAAAAAGGAAAGAGCGAGGCTT
GGGCGGCGATGGAAGCAATTCAGCGCATCGTGCCGTGGGATTTAGCAGCAACCATCATAGT
TGAAGACGAAGAAATAGCTAAAAAAATAAAAGAAATTATTAAGAAAAAAGTGCCCGAGGCTTA
CATTGTGGTTACAGCAAACATAATAGTGATCAATGCACAAACGGAGGAATTATTAAAAATCGC
ATTAGAAGAAGCAAGAAAAATTTTCCTCGAGCACCACCACCACC

Des747:

CTCTAGAAATAATTTTGTTTAACTTTAAGAAGGAGATATACCATGGGCGACGAGGAGGAGAAA
AAGGCCTTGGAGCTTTTCCAAAAAATCTGGGAGGAAGCTAAAAAAAAAGGGGTCTCGGAGG
AGGAAGCTCTGTGGATGGCAGTGTACTACTTGAAACACCAGGCGAATGCACACATAGTGTT
CCTGTCAGAGGACGAGCGGGAAGCAAAAAAGATTCAAGAGATAATCAAGCGGCTGATCCCA
GAAGCGACTGTACTTACGAATTTCGGCGTTGTGGTAGTCTTGGCCCGGTCGGAAGAGTTAA
AAAAATTAGTATCGCTTCTTCAACGTAAAGCCGTGCTCGAGCACCACCACCACC

BB5523:
GTGACAAGAGAGGAAATCGTTAACCTTACAGTCGAACTTTTCAAGAATAAAGAAAGCCCTAG
AGCCACAGAGTTATTAGAGGAGCTGTCAAAAAAGCTGAAGGAGGCCGGGATCGAGGACTTT
ACGGCGATTGAGTTGGACTATAGCGAAGAGCGTTTGAAAGTCCTGAAGGAACTGGAGAAGA
AAGAGAAAGATATCAGCGTAGTCGAAGTGGACGGCAAAATTATAGTCACCGCTTTTACTAAA
GAGAGCGCGGAAGTTATAGAAAAGACTCTTAAAGAAATCGAG

T42A:

CTCTAGAAATAATTTTGTTTAACTTTAAGAAGGAGATATACCATGGGCGTTACGCGCGAAGAG
ATCGTCAACCTTACCGTAGAGCTGTTTAAGAATAAGGAGTCCCCGCGTGCTACAGAGCTTCT
TGAGGAGTTGAGTAAGAAGTTAAAGGAGGCGGGAATAGAGGATTTCGCCGCTATAGAGTTG
GATTACTCTGAGGAGCGCCTGAAGGTACTGAAAGAGTTAGAAAAGAAGGAGAAAGATATCTC
TGTCGTCGAAGTGGACGGAAAAATCATAGTTACTGCTTTCACAAAAGAGTCTGCTGAGGTCA
TCGAGAAGACATTGAAGGAGATCGAGCTCGAGCACCACCACCACC

T42N:

CTCTAGAAATAATTTTGTTTAACTTTAAGAAGGAGATATACCATGGGCGTAACGCGCGAGGAA
ATCGTGAACCTGACTGTCGAACTGTTCAAGAATAAGGAAAGCCCTCGGGCTACAGAGCTGC
TTGAGGAATTAAGCAAAAAGCTGAAGGAGGCAGGCATCGAAGACTTTAACGCCATCGAACT

TGATTACTCTGAAGAACGCCTGAAGGTCCTGAAGGAGCTTGAGAAGAAGGAAAAAGATATAT
CTGTCGTAGAGGTTGATGGCAAGATCATTGTGACTGCCTTTACAAAGGAATCTGCGGAAGTG
ATCGAGAAGACTCTTAAGGAAATCGAACTCGAGCACCACCACCACC

I44A:

CTCTAGAAATAATTTTGTTTAACTTTAAGAAGGAGATATACCATGGGCGTTACGCGGGAGGAA
ATCGTGAACTTGACAGTTGAGCTGTTTAAAAACAAGGAGAGTCCTCGTGCGACCGAGTTAC
TGGAAGAACTGTCTAAGAAACTTAAGGAAGCCGGTATAGAGGACTTCACAGCTGCTGAATTG
GATTACTCTGAGGAACGGTTGAAAGTACTTAAGGAGTTAGAGAAAAAAGAGAAGGACATATC
CGTGGTGGAGGTAGACGGCAAGATCATCGTCACCGCGTTTACCAAGGAGTCAGCTGAAGTT
ATCGAGAAAACTCTGAAAGAGATAGAGCTCGAGCACCACCACCACC

L46A:

CTCTAGAAATAATTTTGTTTAACTTTAAGAAGGAGATATACCATGGGCGTGACGAGAGAAGAG
ATTGTCAATTTAACTGTAGAGTTATTCAAGAATAAAGAGAGTCCCCGCGCGACAGAACTTTTA
GAAGAGCTGTCCAAGAAGTTGAAGGAAGCGGGGATTGAAGACTTTACTGCAATAGAGCTGG
ACTACTCCGAAGAACGGCTGAAGGTTCTTAAGGAGTTAGAAAAGAAGGAAAAGGATATATCC
GTGGTCGAAGTGGACGGCAAGATCATAGTCACAGCCTTCACAAAGGAGTCCGCCGAGGTA
ATTGAGAAAACCTTAAAGGAAATTGCCCTCGAGCACCACCACCACC

T42A/I44A/L46A:

CTCTAGAAATAATTTTGTTTAACTTTAAGAAGGAGATATACCATGGGCGTCACACGGGAGGAG
ATTGTTAATTTAACCGTCGAATTATTTAAAAATAAGGAAAGCCCCCGGGCCACCGAGCTTCTT
GAAGAGTTGTCGAAGAAACTTAAGGAGGCCGGAATCGAAGACTTCGCAGCAGCGGAGGCC
GATTACAGCGAAGAGCGCTTAAAAGTCCTGAAAGAGCTGGAAAAGAAAGAAAAAGATATCTC
CGTAGTCGAGGTGGATGGCAAAATTATCGTTACAGCATTTACAAAGGAGTCCGCAGAAGTCA
TCGAGAAGACCTTGAAAGAGATAGAACTCGAGCACCACCACCACC
